# 4D DYNAMIC SPATIAL BRAIN NETWORKS AT REST LINKED TO COGNITION SHOW ATYPICAL VARIABILITY AND COUPLING IN SCHIZOPHRENIA

**DOI:** 10.1101/2023.09.18.558295

**Authors:** Krishna Pusuluri, Zening Fu, Robyn Miller, Godfrey Pearlson, Peter Kochunov, Theo G.M. Van Erp, Armin Iraji, Vince D. Calhoun

**Affiliations:** Tri-institutional Center for Translational Research in Neuroimaging and Data Science (TReNDS), Georgia State University, Georgia Institute of Technology, and Emory University 55 Park Pl NE, Atlanta, GA 30303, USA; Department of Computer Science, Georgia State University, Atlanta, GA 30303, USA; Department of Psychiatry, Yale School of Medicine, New Haven, CT 06511, USA; Department of Psychiatry, University of Maryland School of Medicine, Baltimore MD 21201, USA; Clinical Translational Neuroscience Laboratory, Department of Psychiatry and Human Behavior University of California Irvine, Irvine Hall, room 109, Irvine, CA, 92697-3950, USA; Center for the Neurobiology of Learning and Memory, University of California Irvine 309 Qureshey Research Lab, Irvine, CA, 92697, USA

**Keywords:** Resting state fMRI (rsfMRI), spatial network dynamics, brain networks, inter-network dynamic volumetric coupling, schizophrenia

## Abstract

Despite increasing interest in the dynamics of functional brain networks, most studies focus on the changing relationships over time between spatially static networks or regions. Here we propose an approach to study dynamic spatial brain networks in human resting state functional magnetic resonance imaging (rsfMRI) data and evaluate the temporal changes in the volumes of these 4D networks. Our results show significant volumetric coupling (i.e., synchronized shrinkage and growth) between networks during the scan. We find that several features of such dynamic spatial brain networks are associated with cognition, with higher dynamic variability in these networks and higher volumetric coupling between network pairs positively associated with cognitive performance. We show that these networks are modulated differently in individuals with schizophrenia versus typical controls, resulting in network growth or shrinkage, as well as altered focus of activity within a network. Schizophrenia also shows lower spatial dynamical variability in several networks, and lower volumetric coupling between pairs of networks, thus upholding the role of dynamic spatial brain networks in cognitive impairment seen in schizophrenia. Our data show evidence for the importance of studying the typically overlooked voxelwise changes within and between brain networks.

## 1. INTRODUCTION

Resting-state functional magnetic resonance imaging (rsfMRI) investigates spontaneous neural activity indirectly via bloodoxygen-level-dependent (BOLD) signal [1,2]. Human rsfMRI data have been extensively studied to identify brain networks with coherent patterns of spatio-temporal activity that are disrupted in numerous brain disorders including schizophrenia (SZ) [3–8]. Recent studies examined temporal fluctuations by employing sliding-window approaches to measure dynamic functional network connectivity (dFNC) and found that SZ is strongly associated with the temporal dynamics of whole-brain network connectivity [4, 9, 10]. A common assumption for most studies is that these networks are fixed spatially over the course of a typical scan, failing to fully capture the highly dynamic nature of the brain networks whose functional connectivity can vary over time, while the networks themselves can also evolve temporally at the voxel-wise scale (e.g., shrink or grow) [11]. A model of the spatial chronnectome was proposed in [5] to identify spatially dynamic network features and voxel-level variations in network coupling that were affected in SZ. Cognitive performance, symptom severity, and drug scores of subjects and their links to resting state dynamic functional network connectivity, multimodal biomarkers, and multi-spatial-scale dynamic functional connectivity have been previously investigated [12–15]. Data-driven analytical approaches such as independent component analysis (ICA) typically assume linear separability of spatial and temporal dynamics of brain networks. Here we introduce an approach to study spatially dynamic 4D brain networks (3D voxel-wise changes over time windows) by utilizing a volumetric measure. We show that significant volumetric coupling (i.e., synchronized shrinkage and growth) exists between networks over time. In addition, the spatial variability of these dynamic spatial brain networks and the coupling between network pairs are modulated differently in SZ patients compared to typical controls. Linear regression analysis reveals that dynamic spatial brain network features are associated with cognitive performance, drug dosage and symptom scores. This work highlights the importance of studying the (voxel-wise) spatial dynamics of brain networks [5] that could characterize brain health and disorder, and help in the development of relevant biomarkers.

## 2. METHODS

### 2.1 Data Collection and Preprocessing

We analyzed 3-Tesla rsfMRI data from 508 subjects collected in three studies The Functional Imaging Biomedical Informatics Research Network (FBIRN) [4], the Center for Biomedical Research Excellence (COBRE) [6], and a Maryland Psychiatric Research Center (MPRC) [7] study. In addition to the previously described inclusion criteria [16], we only selected the 508 subjects who also have genomic data (for use in other parts of this project), resulting in 315 controls (CN) and 193 SZ patients. rsfMRI data preprocessing was performed using the statistical parametric mapping toolbox (SPM12, http://www.fil.ion.ucl.ac.uk/spm/). Further details of the datasets and preprocessing can be found in [16].

### 2.2 Analysis pipeline

The analysis pipeline is depicted in Fig. 1. In the first step, we performed group-level spatially constrained independent component analysis (sICA) of rsfMRI data with 20 components, as described in [5], using the group ICA of fMRI toolbox (GIFT) software package [17] (https://trendscenter.org/software/gift/). Of these 20 components, 14 were identified as neuro-related brain networks, as shown in Fig. 2. sICA identifies and separates these brain intrinsic connectivity networks (ICNs) and their associated time courses. In the second step, a sliding-window approach combined with spatially constrained ICA was performed for each subject across time windows, using multi-objective optimization ICA with reference (MOO-ICAR) [18, 19]. This ensures the correspondence of ICNs across subjects and time windows by using the components obtained from group level ICA analysis in the previous step as reference. Our recent study showed that MOO-ICAR is capable of effectively estimating large-scale networks from short time segments [20]. This step captures, for each subject, the brain networks (ICNs) that are spatially varying over time (windows), as well as their time courses within each window. This allows for further analysis of such spatially dynamic brain networks, inter-network couplings, as well as different modulation under disease conditions. A sliding window length of 60s (30 × *TR*, where the repetition time *TR* = 2*s*) was employed based on previous research [5] and falling within the recommended ranges from multiple studies [17]. In the third step, we perform a volumetric analysis for each subject by measuring the volume of each dynamic spatial brain network (computed as the number of z-scored voxels with activity above a volume threshold zscore, *V*_*th*_, picked from the values [0.5, 1, 1.5, 2, 2.5, 3, 3.5]) as a function of time (windows). We also compute the volumetric coupling (correlations converted into Fisher’s z-scores) between such networks over time, resulting in a 14x14 volumetric coupling (VC) matrix for each subject, corresponding to the 14 relevant brain networks, as shown in Step 3 of Fig. 1. A large positive coupling/correlation value in the matrix implies that the two corresponding networks grow and shrink together, while a negative value implies that one network grows while the other shrinks. In the fourth step, we use the subject-level VC matrices, as well as the mean and standard deviations (std) of the volumes of networks, to perform hypothesis testing. We identify dynamic spatial brain networks with significant VC in each group using 1-sample t-tests, as well as study group differences in mean/std network volumes and VC of network pairs between the control group and individuals with schizophrenia using 2-sample t-tests. A 5% false discovery rate (FDR) [21] is applied to correct for multiple comparisons at each threshold or across all thresholds (results shown in bold). Performing such analyses at multiple volume thresholds (*V*_*th*_) allows us to examine the networks at different levels of granularity, and identify changes in regions of either more focused or wide spread activity. We also perform robust linear regression analysis of cognitive scores of all subjects, and symptom and drug scores of patients against the mean/std network volumes and VC of network pairs as predictors, while using age, gender and scanning site as covariates. The positive and negative syndrome scale (PANSS) scores [13] are available for the FBIRN and COBRE datasets, while Brief Psychiatric Rating Scale (BPRS) scores are available for MPRC dataset, which are then converted into PANSS total scores using the matching obtained in [15,22]. Chlorpromazine (CPZ) equivalent drug scores [13,23] are available for all three datasets. Across both patients and controls, we also employed two cognitive scores [14] (processing speed, working memory) available for all three datasets and two cognitive scores (visual learning, verbal learning) available only for FBIRN and COBRE, after separately normalizing the scores for each dataset. On the whole, 178 patients had PANSS total scores, 111 patients had CPZ drug scores, 459 subjects had processing speed scores, 377 subjects had working memory scores, 254 subjects had verbal learning scores, and 251 subjects had visual learning scores.

**Fig. 1.**
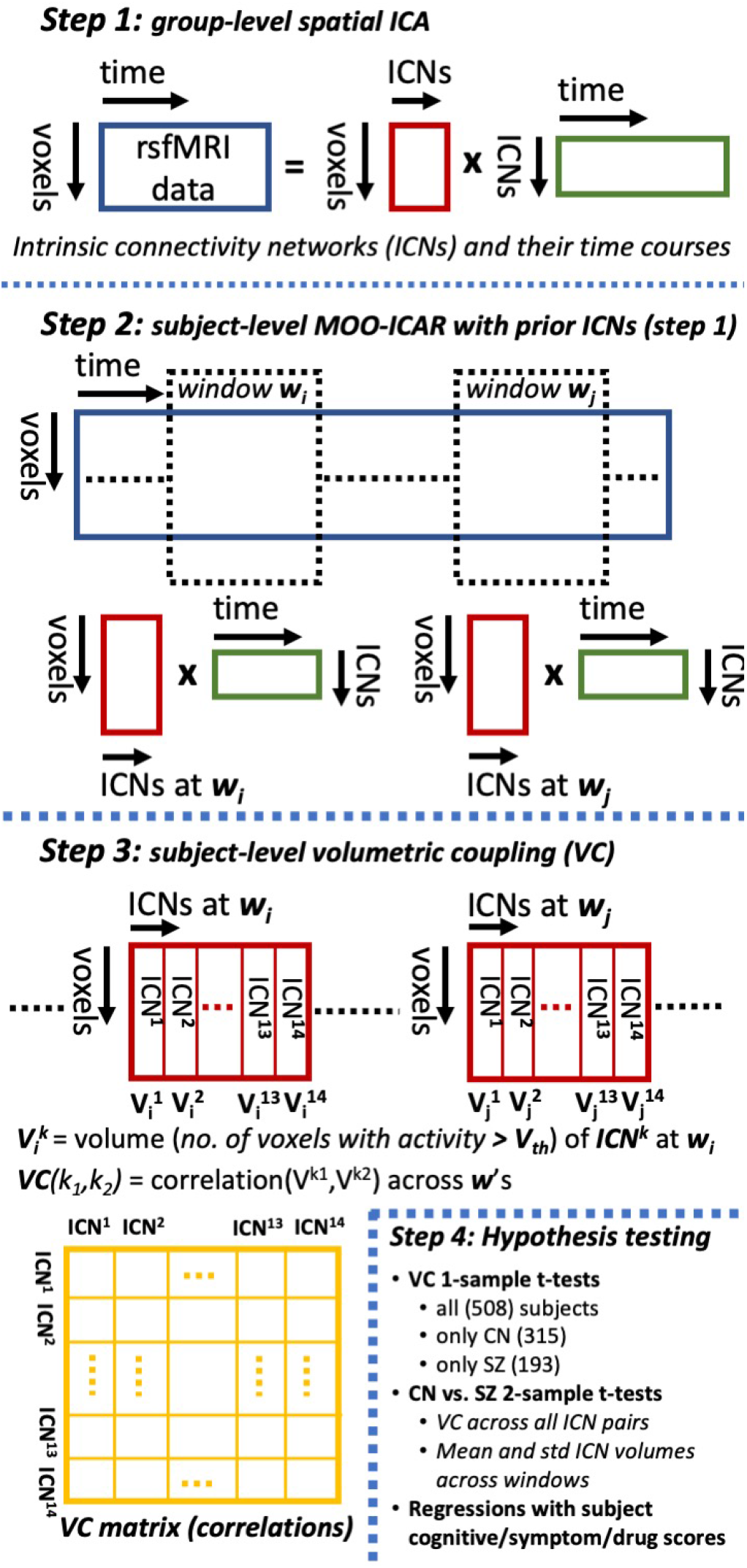
Analysis pipeline showing **(1)** group level spatial ICA followed by **(2)** subject level MOO-ICAR over sliding windows using prior components/spatial networks (ICNs) from step (1) as reference to maintain correspondence of ICNs across subjects and time windows. **(3)** Volume of each network at each window is computed as the number of voxels with z-scored activity above a volume threshold, *V*_*th*_. Volumetric coupling (VC) matrices are computed at the subject level (measured as correlations between volumes of ICNs across time windows). Further analysis is done using **(4)** 1sample, 2-sample t-tests and regression analysis. Detailed description of the pipeline is presented in Section 2.2.

**Fig. 2.**
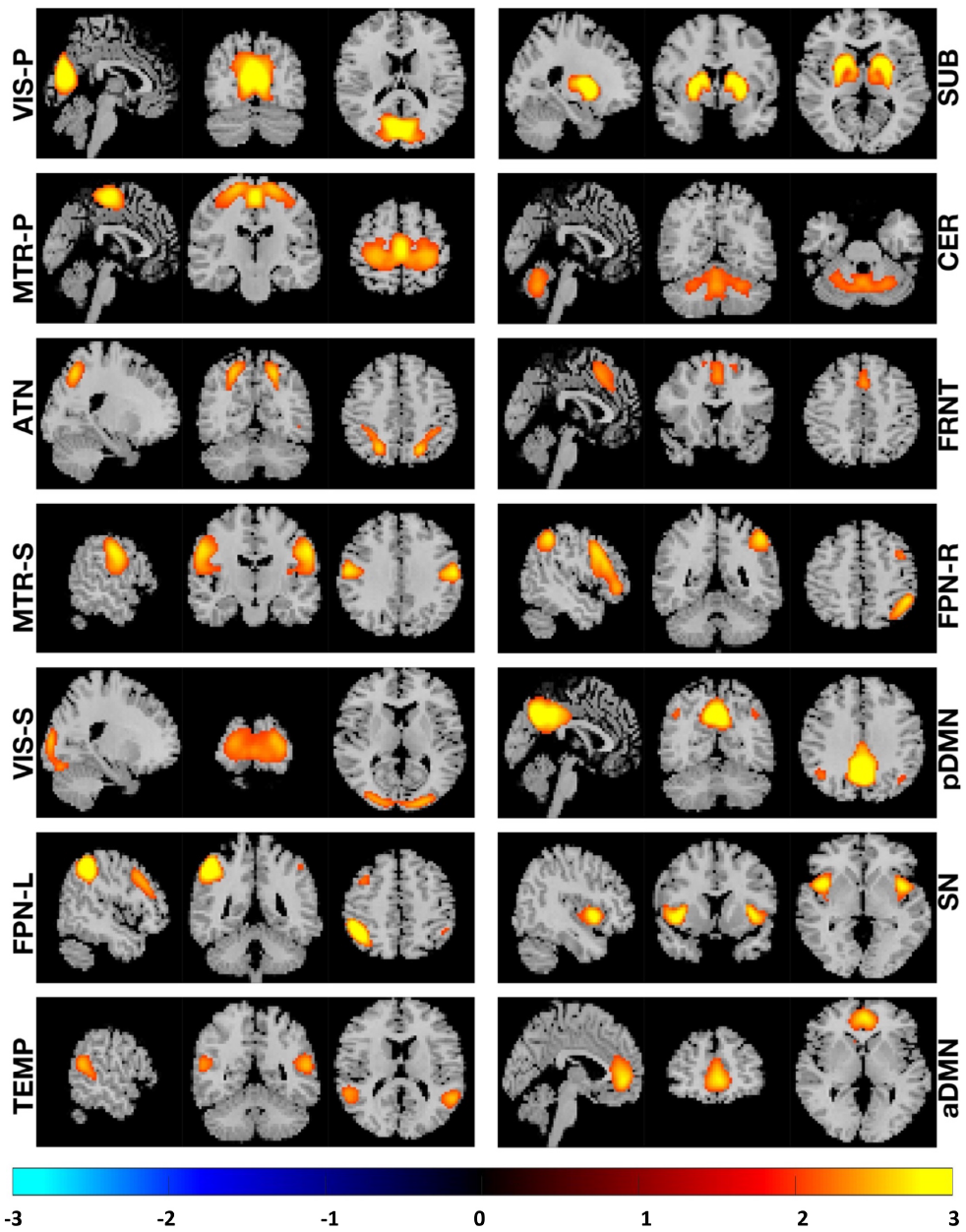
The mean activity maps across typical control subjects for each of 14 relevant brain networks are shown along three planar cross sections sagittal, coronal, and transverse taken near the voxel with the highest activity. The maps are z-scored and only show regions with *z ≥*2, with anatomical images overlaid in the background. *V IS −P* (visual primary), *SUB* (subcortical), *MTR −P* (somatomotor primary), *CER* (cerebellar), *ATN* (attention dorsal), *FRNT* (frontal), *MTR −S* (somatomotor secondary), *FPN− R* (frontoparietal right), *V IS S* (visual secondary), *pDMN* (default mode posterior), *FPN −L* (frontoparietal left), *SN* (salience), *TEMP* (temporal), and *aDMN* (default mode anterior).

## 3. RESULTS AND DISCUSSION

### 3.1 Dynamic spatial brain networks

14 relevant brain networks are identified, as shown in Fig. 2. These are labeled *V IS −P* (visual primary), *SUB* (subcortical), *MTR −P* (somatomotor primary), *CER* (cerebellar), *ATN* (attention dorsal), *FRNT* (frontal), *MTR S* (somatomotor secondary), *FPN −R* (frontoparietal right), *V IS −S* (visual secondary), *pDMN* (default mode posterior), *FPN −L* (frontoparietal left), *SN* (salience), *TEMP* (temporal), and *aDMN* (default mode anterior) networks. These networks are spatially dynamic at the voxel level and undergo changes in volumes over time, as demonstrated in Fig. 3 for the networks *CER* and *pDMN* in CN group. The figure shows the mean activity maps and demonstrates the variability in the volumes of these networks across CN subjects over windows with the largest volumes (top 10%) or the smallest volumes (bottom 10%), measured at *V*_*th*_ = 2.

**Fig. 3.**
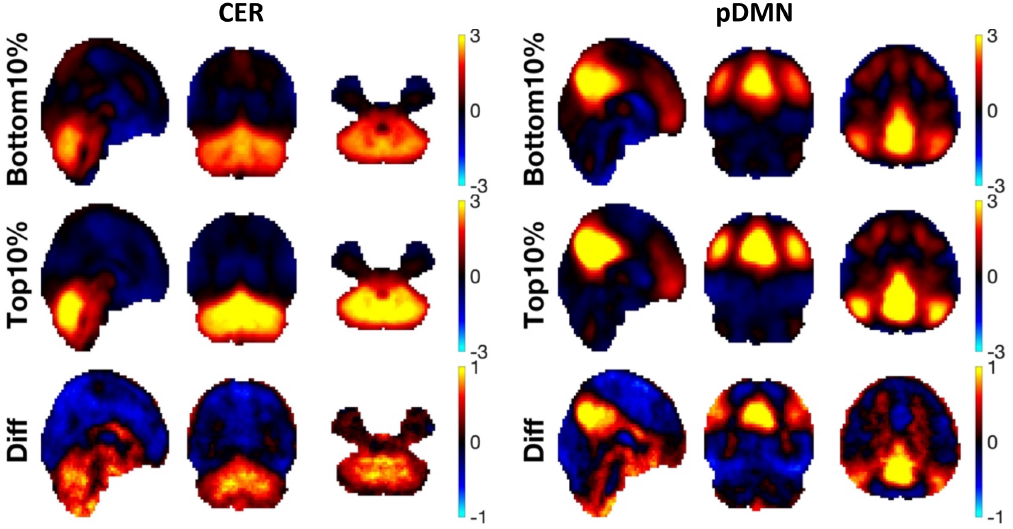
Spatial variability and regions with volumetric changes in cerebellar (*CER* – left) and posterior default mode (*pDMN* – right) networks revealed using mean activity maps across subjects in CN group, over windows with bottom 10% smallest volumes or top 10% largest volumes measured at *V*_*th*_ = 2, and the differences between the two.

In addition, the difference maps between the largest and the smallest mean volumes depict the regions where volumetric changes occur. As can be inferred from Figs. 2 & 3 and the color bars next to them, by only examining regions above a particular *V*_*th*_, one can investigate either the most active/focused region of the network (high *V*_*th*_) or a more widespread region (low *V*_*th*_).

### 3.2 Altered focus of activity and lower spatial network dynamics in SZ

Volume of each network was measured per subject per window at different volume thresholds (*V*_*th*_) to determine subject level mean and standard deviations (std) of network volume. We analyze results at different volume thresholds to determine whether the most active regions of the network are affected differently from the wider network. Two sample t-test results comparing CN and SZ groups for mean volumes of networks are shown in Table 1, while those for the std volumes using non-parametric rank-sum tests are shown in Table 2, with a 5% false discovery rate (FDR) [21] for multiple comparisons at each threshold or across all thresholds (in bold). Table 1 reveals that several networks show significant mean volume changes in SZ as they grow or shrink during the scan. For *SUB, CER* and *FPN −R* networks, CN shows lower volumes (blue) at low thresholds and higher volumes (red) at higher thresholds compared to SZ, implying that more focused activity in these networks is lowered in SZ as activity becomes widespread. For the networks *V IS −S* and *TEMP*, it is vice versa with activity becoming more focused and less widespread in SZ. *ATN* and *pDMN* networks only shrink at moderate thresholds. Table 2 shows that in the case of std of volumes, the networks *V IS −P, MTR −P, CER, ATN, FRNT, MTR S, V IS− S, pDMN* and *TEMP* have lower variability or spatial dynamics in SZ at different thresholds, while no networks seem to have significantly higher variability in SZ as compared to CN. For each network, results at different thresholds reveal how focused or widespread are the reduced spatial dynamics seen in SZ. For example, the network *MTR −S* shows most significantly lower spatial dynamics in SZ at *V*_*th*_ = 1, which gradually become less significant as we move towards more widespread (smaller *V*_*th*_) or more focused (larger *V*_*th*_) regions of the network.

**Table 1.**
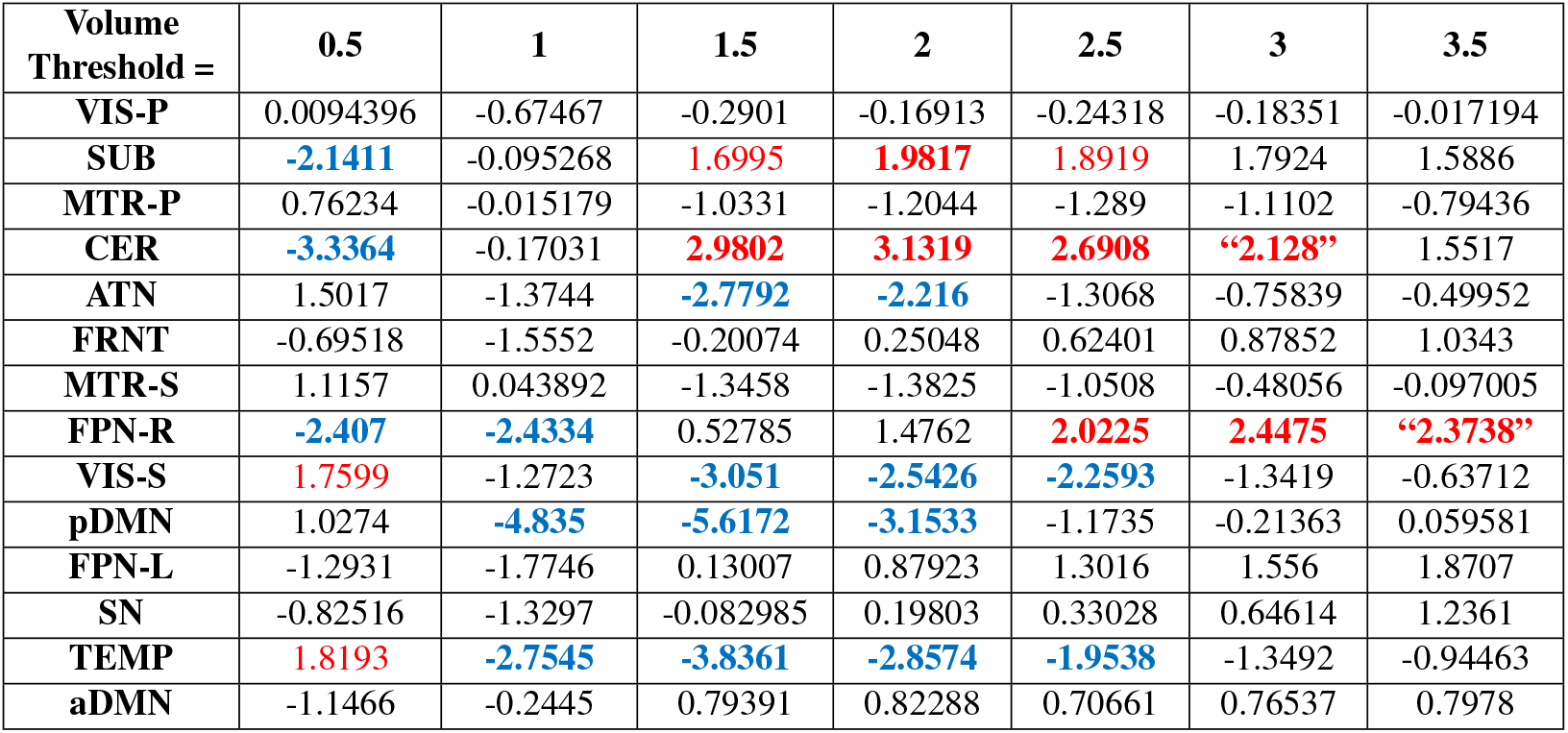
(*CN −SZ*) two-sample t-test results comparing mean network volumes across CN and SZ groups at different volume thresholds. *k* =*− log*10(*p*) *∗sign*(*t*) values are shown, with significant ones depicted in red for *CN > SZ*, and blue for *CN < SZ*. Significant results at each threshold with 5% FDR for multiple comparisons (14 comparisons, one for each network) are shown, while those that remain significant with 5% FDR across all thresholds (14 *×*7 comparisons) are shown in bold. Note that *CER* network at *V*_*th*_ = 3, and *FPN− R* network at *V*_*th*_ = 3.5 (shown in double quotes) are significant with FDR across all thresholds, but not with FDR at single threshold.

**Table 2.** (*CN− SZ*) 2-sample non-parametric rank-sum test results comparing the standard deviation of network volumes over windows per subject across CN and SZ groups at different volume thresholds. *k* =*− log*10(*p*) *∗ sign*(*z*) values are shown, with significant *CN > SZ* ones depicted in red (no networks found with significant *CN < SZ*). Significant results at each threshold with 5% FDR for multiple comparisons (14 comparisons, one for each network) are shown, while those that remain significant with 5% FDR across all thresholds (14 *×*7 comparisons) are shown in bold. Note that *CER* network at *V*_*th*_ = 2.5, 3 (shown in double quotes) is significant with FDR across all thresholds, but not with FDR at single thresholds.

| Volume Threshold = | 0.5 | 1 | 1.5 | 2 | 2.5 | 3 | 3.5 |
| --- | --- | --- | --- | --- | --- | --- | --- |
| VIS-P | 0.90706 | **1.8232** | **1.6755** | 1.4843 | 0.24438 | -0.39964 | -0.80985 |
| SUB | 0.70326 | 1.0632 | 1.1278 | 0.60998 | 1.2882 | 1.2383 | 0.99868 |
| MTR-P | **1.9253** | **2.1333** | **3.2954** | **2.8933** | 0.24956 | -0.44333 | -1.073 |
| CER | 1.5615 | 0.98867 | 1.4843 | 1.2669 | **“2.2754”** | **“2.0787”** | 1.4702 |
| ATN | **2.2537** | **2.3241** | **2.7664** | **2.0437** | -0.079203 | -1.1611 | -1.1815 |
| FRNT | 0.36165 | 1.1527 | **1.9253** | 0.19987 | -0.25118 | 0.20323 | 0.56631 |
| MTR-S | **2.0564** | **3.702** | **3.4156** | **2.9502** | 1.0961 | -0.29175 | -1.0386 |
| FPN-R | 1.1521 | 0.76978 | 1.4329 | 1.4521 | 1.5733 | 1.207 | 1.0758 |
| VIS-S | 0.71425 | 1.592 | **2.0414** | 1.4924 | 0.14984 | -0.84875 | -1.618 |
| pDMN | 0.24085 | 1.3512 | **2.675** | 0.15411 | -0.79823 | -0.49077 | -0.25248 |
| FPN-L | 1.2426 | 0.44649 | 1.3667 | 1.1967 | 0.76978 | 0.74373 | 1.2266 |
| SN | 0.28565 | 1.5809 | 1.0004 | 0.67029 | 0.6536 | 0.46802 | 0.52484 |
| TEMP | **2.554** | **2.2529** | **3.0793** | **1.9662** | 0.064951 | -0.98922 | -1.2315 |
| aDMN | 0.7902 | 0.59129 | 1.4012 | 0.98978 | 0.93498 | 0.49941 | 0.36678 |

### 3.3 Dynamic networks show significant volumetric coupling

Volumetric coupling between spatial dynamic network pairs was measured per subject as the correlation between their volumes over time windows, measured at different volume thresholds (*V*_*th*_). Results from one sample t-tests checking for non-zero correlations, as shown in Fig. 4 and Table 3, demonstrate that several dynamic spatial network pairs across groups have significant VC. Positive VC between two networks indicates that both networks grow and shrink together, while negative VC indicates one network grows while the other shrinks. The connectogram in Fig. 4 at *V*_*th*_ = 2 shows strong positive coupling (yellow/orange lines) between the network pairs (*V IS −P/V IS −S*), (*MTR −P/MTR −S*) and (*FPN−R/FPN −L*). The network *aDMN* has a strong negative coupling (blue line) with *ATN*, and a strong positive coupling (red line) with *FRNT* . Similarly, the network SN has strong negative coupling (blue line) with *pDMN*, and strong positive coupling (red lines) with both the networks *SUB* and *MTR S. −*A false discovery rate (FDR) of 5% to correct for multiple comparisons [21] at this threshold shows that out of 91 possible network pair combinations for the 14 relevant brain networks, 62 network pairs in controls have significant VC, of which 34 are positively, and 28 are negatively coupled. A similar analysis for patients with SZ shows that 45 brain network pairs have significant VC, with 30 network pairs having positive and 15 networks having negative coupling. For the combined dataset with subjects from both groups, 65 network pairs have significant VC, with 35 positively and 30 negatively coupled. The total number of network pairs with significant +ve/-ve VC at different volume thresholds are depicted in Table 3, which shows that several network pairs are volumetrically coupled across groups with 5% FDR at threshold-level, and most of the network pairs show significant coupling even with 5% FDR applied across all thresholds (in bold).

**Table 3.**
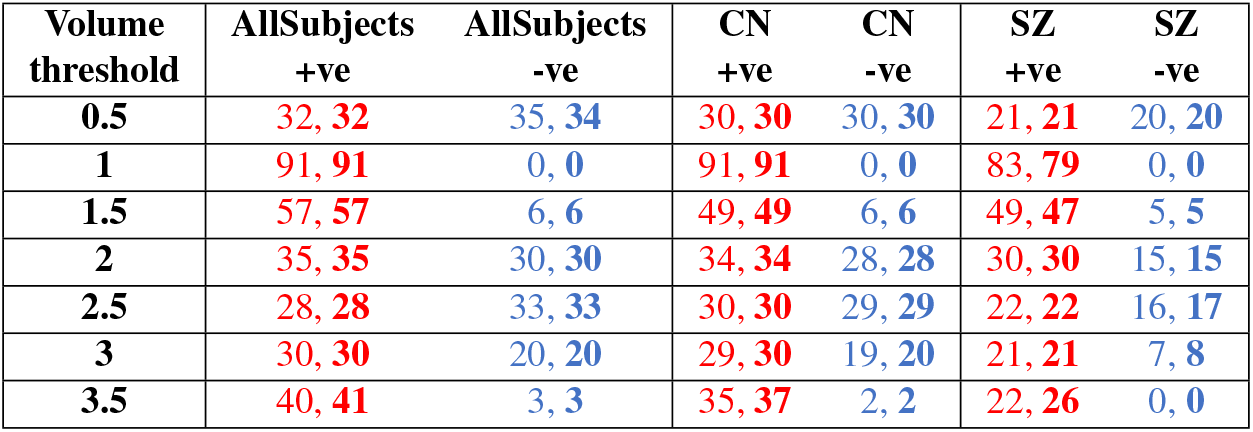
Number of network pairs with significant non-zero +ve (red) or -ve (blue) volumetric coupling as found from 1-sample t-test results at different volume thresholds (out of 91 possible network pair combinations, chosen from 14 networks). Results with 5% FDR correction applied for multiple (91) comparisons at each given threshold are shown, while significant results with 5% FDR correction applied across all thresholds (91 *×*7 total comparisons) are shown in bold. The CN group consistently shows more network pairs with significant +ve/-ve volumetric coupling than the SZ group.

**Fig. 4.**
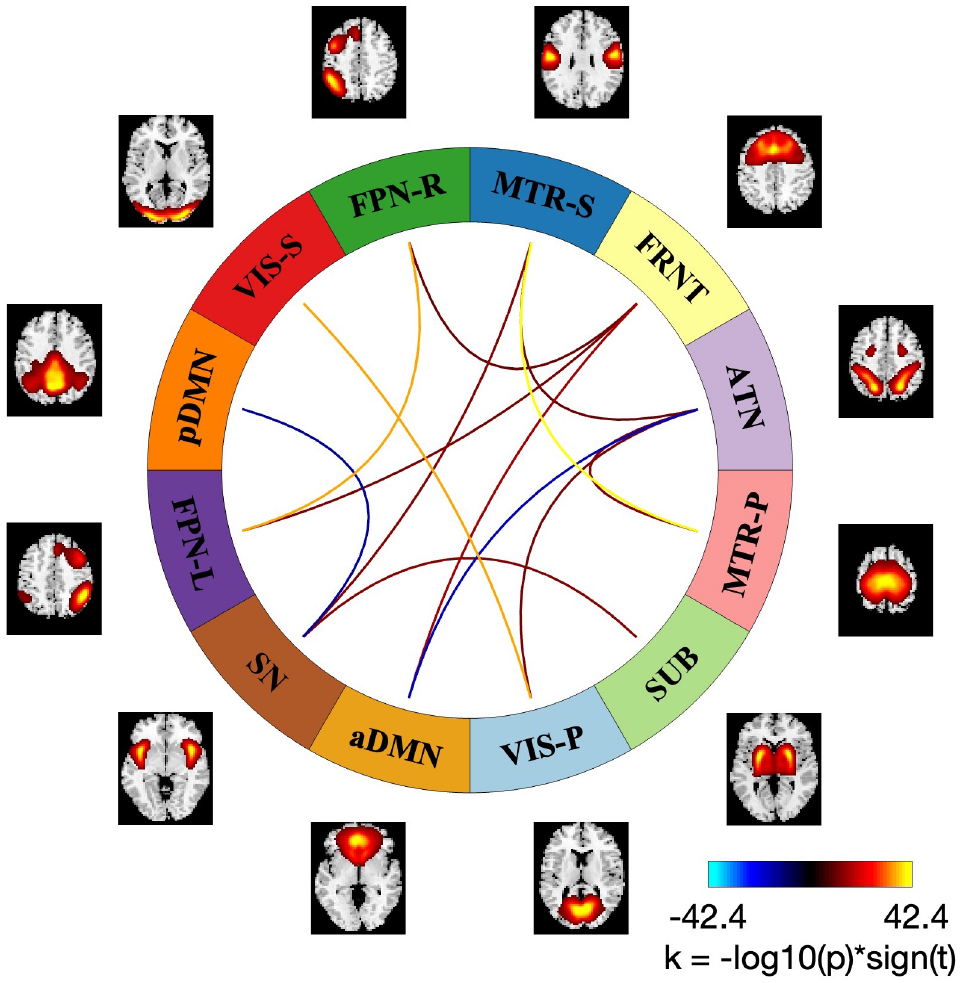
A connectogram depicting networks with strong volumetric coupling, based on the results of one sample t-tests for controls at *V*_*th*_ = 2. Connections are drawn using the values of *k* =*− log*10(*p*) *∗sign*(*t*) for each network pair, where *t* is the computed t-statistic, and *p* is the resulting p-value. Only strong connections with *abs*(*k*) *>* 10 are shown, while the 5% FDR threshold for multiple comparisons at *abs*(*k*) = 1.5391 (corresponding to the critical p-value of 0.0289) shows that 62 network pairs (out of 91 possible combinations) have significant volumetric coupling, of which 34 are positively coupled (i.e., when one network grows, so does the other) and 28 negatively coupled (i.e., when one network grows, the other shrinks). The connectogram shows strong positive coupling (yellow/orange lines) between the network pairs (*V IS − P/V IS − S*), (*MTR − P/MTR − S*), (*FPN − R/FPN − L*), while strong negative coupling (blue lines) exists between the networks (*aDMN/ATN*), (*pDMN/SN*).

### 3.4 Reduced volumetric coupling in SZ

One-sample t-test results in Table 3 show that fewer number of dynamic spatial brain networks show significant VC in SZ compared to CN group, implying that VC, whether positive or negative, is lower in SZ as compared to CN. Two-sample ttests of subject level VC measures were performed at different *V*_*th*_ to identify volumetrically coupled network pairs with different modulations under SZ disease conditions as compared to CN. Multiple comparisons were accounted for using 5% FDR correction at each threshold or across all thresholds (in bold). As shown in Table 4, we identified several network pairs with significant group differences in (*CN − SZ*) 2sample t-tests. A significantly negative (*CN− SZ*) t-statistic could imply that either SZ patients gained positive VC that didn’t exist in CN, or that SZ patients lost negative VC that existed in CN. Similarly, a significant positive (*CN− SZ*) t-statistic could imply either that SZ patients lost positive VC existing in CN, or that SZ patients gained -ve VC that didn’t exist in CN. This is resolved by looking at the corresponding 1-sample t-test results in Table 4, which show that in all the cases SZ patients lost either positive/negative VC existing in CN, but no networks gain positive/negative VC in SZ compared to CN. The network pairs (*V IS −S/FPN −R*) at *V*_*th*_ = *{*0.5, 2, 2.5 *}*and (*SUB/V IS − P*) at *V*_*th*_ = *{*0.5, 2*}* have significant negative VC in CN, which is lost in SZ. The network pairs (*TEMP/MTR −S*), (*CER/SUB*) at *V*_*th*_ = 0.5 and (*SN/V IS −S*) at *V*_*th*_ = 1 have significant positive VC in CN, which is lost in SZ. The network pair (*aDMN − FRNT*) at *V*_*th*_ = 1 shows highly significant positive VC in CN, that is somewhat lost in SZ as shown by 2-sample test results, but still remains significant in one-sample t-test results. Even with FDR applied across all thresholds, the positive coupling of (*SN/V IS −S*) at *V*_*th*_ = 1 and the negative coupling of (*V IS S/FPN −R*) at *V*_*th*_ = 2 remain significantly higher in CN compared to SZ.

**Table 4.** Summary table showing dynamic spatial brain network pairs whose volumetric coupling is significantly affected under SZ; Significant (*CN −SZ*) 2-sample t-test results with 5% FDR correction for multiple (91) comparisons at each *V*_*th*_ are shown, along with the corresponding 1-sample t-test results; All values shown are *k* = *−log*10(*p*) *∗sign*(*t*). Network pairs shown in blue lost -ve VC in SZ compared to CN group, those shown in red lost +ve VC in SZ, while the one shown in orange lost some +ve VC in SZ compared to CN in 2-sample t-test, but still remains significant in one sample t-test in SZ; Results that remain significant with 5% FDR across all thresholds (91 *×* 7 comparisons) are shown in bold (*SN/V IS − S* at *V*_*th*_ = 1 and *V IS − S/FPN − R* at *V*_*th*_ = 2). Results show SZ patients lost +ve/-ve VC compared to the control group (CN).

| Volume Threshold | Network1 | Network2 | 2 Sample t-test (CN-SZ) | 1 Sample t-test (All subjects) | 1 Sample t-test (CN) | 1 sample t-test (SZ) |
| --- | --- | --- | --- | --- | --- | --- |
| 0.5 | SUB | VIS-P | -3.2803 | -5.9809 | -8.4771 | -0.13055 |
|  | VIS-S | FPN-R | -3.538 | -2.6627 | -5.9664 | 0.41982 |
|  | TEMP | MTR-S | 3.3503 | 1.6406 | 4.0571 | -0.76595 |
|  | CER | SUB | 2.9146 | 5.9675 | 7.7499 | 0.22573 |
| 1 | SN | VIS-S | **4.2897** | 12.1594 | 15.1279 | 0.76533 |
|  | aDMN | FRNT | 3.1873 | 25.3679 | 21.6856 | 5.1741 |
| 2 | SUB | VIS-P | -3.3981 | -4.8994 | -7.8222 | 0.019538 |
|  | VIS-S | FPN-R | **-4.0738** | -2.1391 | -5.3376 | 0.79012 |
| 2.5 | VIS-S | FPN-R | -3.3537 | -2.4936 | -5.5555 | 0.41586 |

### 3.5 Association of 4D dynamic spatial brain networks with subject cognitive, symptom and drug scores

In order to investigate possible links between dynamic spatial brain networks and subject cognitive, symptom and drug scores, we performed robust linear regression analysis with 4 cognitive scores (visual learning, verbal learning, processing speed and working memory) for all subjects, chlorpromazine (CPZ) drug score for patients, and the positive and negative syndrome scale (PANSS) total symptom score for patients, each separately regressed against the subject-wise means and standard deviations (std.) of the volumes of 14 different networks across windows, as well as the subject-wise volumetric coupling between network pairs (91 combinations) as predictors. This was repeated at 7 different volume thresholds, for a total of 4998 separate regressions (6 *scores ×*7 *thresholds×* (14 *mean network volumes* + 14 *std. network volumes* + 91 *V C of network pairs*)). In each case, age, gender and imaging site were used as covariates. Table 5 shows the features of dynamic spatial brain networks with significant positive (red) or negative (blue) association to subject scores, with 5% FDR correction applied for multiple comparisons at each given threshold (14 total comparisons for mean/std. and 91 for VC), or 5% FDR correction across all thresholds (14*×* 7 comparisons for mean/std and 91*×* 7 for VC; shown in bold). Results reveal strong association of dynamic spatial brain network features with cognitive, drug and symptom scores.

**Table 5.** Association of dynamic spatial brain networks with subject cognitive, symptom and drug scores via robust linear regression analysis: 4 cognitive scores (visual learning, verbal learning, processing speed and working memory) for all subjects, chlorpromazine (CPZ) equivalent drug score for patients, and the positive and negative syndrome scale (PANSS) total symptom score for patients are each separately regressed against the subject-wise means and standard deviations (std.) of the volumes of 14 different networks across windows, as well as the volumetric coupling between network pairs (91 combinations) aspredictors. This is repeated at 7 different volume thresholds. Age, gender and imaging site are used as covariates in each case. Values shown are *k* = *−log*10(*p*) *∗ sign*(*t*), where *t* is the ratio of the regression coefficient to the standard error of its estimate. Predictors with positive coefficients are shown in red, while those with negative coefficients are shown in blue. Significant results with 5% FDR correction applied for multiple comparisons at each given threshold or across all thresholds (in bold) are shown. Results show strong association of dynamic spatial brain network features with cognitive, drug and symptom scores.

| Score | Vth | Mean network volume |  | Std. network volume |  | Network pairs VC |
| --- | --- | --- | --- | --- | --- | --- |
| Visual learning | 0.5 | SUB (-3.7497)<br>FPN-R (-2.8957)<br>TEMP (3.3449) | CER (-2.7101)<br>FPN-L (-2.1757) |  |  |  |
|  | 1 | SUB (-2.2909)<br>pDMN (-3.7004)<br>TEMP (-2.5697) | VIS-S (-2.0565)<br>SN (-4.0015) | FPN-L (2.219)<br>TEMP (4.4672) |  | VIS-S_SN (3.6095) |
|  | 1.5 | TEMP (-5.2192) |  | VIS-P (3.0656)<br>aDMN (2.1947) | TEMP (2.2133) |  |
|  | 2 | SUB (2.582)<br>FPN-R (2.0186)<br>TEMP (-4.5004) | CER (1.8409)<br>FPN-L (2.0236) | VIS-P (2.535) |  |  |
|  | 2.5 | SUB (3.0009)<br>FPN-R (2.6259) | CER (2.155)<br>TEMP (-3.3722) |  |  |  |
|  | 3 | SUB (3.1906)<br>FPN-R (3.0619) | CER (2.5547)<br>TEMP (-2.2514) | SUB (3.1232)<br>FPN-R (2.0044)<br>TEMP (-1.9108) | CER (1.9753)<br>FPN-L (2.2775) |  |
|  | 3.5 | SUB (3.1741)<br>FPN-R (2.8418) | CER (2.5964)<br>SN (2.0359) | SUB (2.9359)<br>FPN-R (2.211)<br>SN (2.0853) | CER (2.4758)<br>FPN-L (1.7988) |  |
| Verbal learning | 0.5 | CER (-2.5356)<br>FPN-L (-2.6967) | FPN-R (-2.6899) |  |  |  |
|  | 1 |  |  |  |  | VIS-S_SN (3.309) |
|  | 2 | CER (2.1507)<br>FPN-L (2.599) | FPN-R (2.4745) |  |  |  |
|  | 2.5 | CER (2.0852)<br>FPN-R (2.2112) | MTR-S (-2.8836)<br>FPN-L (2.4156) |  |  |  |
|  | 3 | MTR-S (-3.0627) |  |  |  |  |
|  | 3.5 | MTR-S (-2.5588) | SN (2.4424) | SN (2.6854) |  |  |
| Processing speed | 0.5 | TEMP (2.5217) |  | MTR-S (2.5104) |  |  |
|  | 1 | pDMN (-3.1079) |  | MTR-S (2.783)<br>TEMP (2.8588) | SN (2.0394) | VIS-S_SN (3.3209) |
|  | 1.5 | pDMN (-2.7853)<br>TEMP (-3.1043) |  | MTR-S (2.6377) | VIS-S (2.6716) |  |
|  | 2 | TEMP (-3.3893) |  | MTR-S (3.5447) | VIS-S (2.7952) |  |
|  | 2.5 | TEMP (-3.5632) |  | MTR-S (2.5309) |  |  |
|  | 3 | TEMP (-3.014) |  | SN (2.4798) |  |  |
|  | 3.5 | SN (2.6027) |  | SN (3.1808) |  |  |
| Working memory | 0.5 |  |  |  |  | SUB_FPN-R (3.3628) |
|  | 1 |  |  |  |  | MTR-P_ATN (3.0683)<br>FRNT_FPN-L (3.6719) |
|  | 2.5 |  |  |  |  | SUB_FPN-R (3.4186)<br>FPN-R_VIS-S (-3.0587) |
|  | 3 |  |  |  |  | VIS-P_TEMP (3.0692)<br>SUB_FPN-R (3.1227) |
| CPZ drug | 1.5 | MTR-S (-2.4957) |  | SN (2.7973) |  |  |
| PANSS total | 1.5 | CER (2.7301) |  |  |  |  |
|  | 2 | CER (2.6539) |  |  |  |  |
|  | 2.5 | CER (2.5912) |  |  |  |  |

As seen with 2-sample t-test results comparing CN vs. SZ mean network volumes in Section 3.2 and Table 1, regression results in Table 5 for various scores against the mean network volumes in all subjects also reveal how altered focus of activity affects these scores. For example, visual learning shows -ve association with *SUB, CER* and *FPN −R* at low thresholds (*V*_*th*_ = 0.5), and +ve association with those networks at high thresholds (*V*_*th*_ = 2, 2.5, 3, 3.5), implying that more focused activity in these networks has a +ve effect on the score, while less focused or more wide spread activity has a -ve effect. Similarly, +ve associations are seen for more focused activity in *CER, FPN− R* and *FPN −L* with verbal learning, while more wide spread activity is associated with lower scores. For both visual learning and processing speed, more wide spread activity in *TEMP* is associated with improved scores, whereas more focused activity is linked to lower scores.

Regression results for std in Table 5 show that all the scores predominantly show positive association (red) with std network volumes (with the exception of visual learning with *TEMP* at *V*_*th*_ = 3), implying that higher dynamic variability in these networks is linked to improved scores. Comparing these results against the results from 2-sample rank-sum tests for CN vs. SZ std network volumes in Section 3.2 and Table 2 which showed SZ only reduced dynamic variability in these networks, highlights the role of spatial dynamic variability in cognitive performance and SZ.

As described for 2-sample t-test results for VC in SZ vs CN in Section 3.4 and Table 4, regression results against VC can only be resolved properly by also analyzing the corresponding 1-sample VC results looking for non-zero coupling between these network pairs. A significant positive regression coefficient for a score against the VC of a network pair could imply that the network pair has positive VC, and a higher magnitude of such coupling leads to higher performance/score, or alternatively, that the network has -ve VC and higher magnitude of such coupling leads to lower performance/score. Similarly, a significant -ve regression coefficient implies that the network has -ve VC and higher magnitude of the coupling leads to improved scores, or alternatively, that the network has +ve VC and higher magnitude leads to lower scores. In each case, we examined the corresponding 1sample t-test results (values not shown) for non-zero VC and found that +ve regression coefficients against VC in Table 5 are due to higher magnitudes of +ve VC in the corresponding networks associated with improved scores, and -ve regression coefficients are due to higher magnitudes of -ve VC in the corresponding networks associated with improved scores, thus highlighting in both cases that improved scores are associated with stronger VC (+ve/-ve), whereas SZ is associated with the loss of such VC (+ve/-ve) between network pairs as seen in Section 3.4 and Table 4.

## CONCLUSIONS

We identify several spatially dynamic brain networks that expand or shrink over time. SZ is linked to such network dynamics, as some networks show higher volumes in SZ (*ATN, pDMN*), some show predominantly less focused activity and higher wide spread activity (*SUB, CER, FPN− R*), while a few others show predominantly higher focused activity and lower wide spread activity (*V IS S, TEMP*). In addition, several networks (*V IS −P, MTR −P, CER, ATN, FRNT, MTR −S, V IS −S, pDMN, TEMP*) show lower spatial dynamical variability in SZ compared to CN, while no networks show higher variability in SZ. We also demonstrate that dynamic spatial brain networks show significant volumetric coupling with synchronized shrinkage/expansion. Such network coupling is altered in SZ, and patients have significantly lower positive (*TEMP/MTR −S, CER/SUB, SN/V IS −S, aDMN/FRNT*) or negative (*SUB/V IS −P, V IS −S/FPN −R*) volumetric coupling between network pairs compared to controls. On the other hand, no network pairs were found to have significantly higher positive/negative volumetric coupling in SZ. Robust linear regression analyses show that several features of such dynamic spatial brain networks, such as the subjectwise means or standard deviations of network volumes and the volumetric coupling between network pairs are associated with cognitive performance, CPZ equivalent drug dosage and symptom scores, as well as highlight the role of altered focus of activity in these networks and its effects on these scores. While cognitive, symptom and drug scores show strong +ve association with higher dynamic variability in network volumes and higher VC (+ve/-ve) between network pairs, SZ shows lower dynamic variability in the volumes of these networks as well as lower VC between network pairs compared to controls, suggesting a possible mechanism for cognitive defects of SZ wherein reduced network-level spatial dynamics and reduced inter-network VC could lead to diminished performance.

Future work should replicate these findings in larger datasets and further examine the dysfunction of the visual system [24, 25], the reduced suppression of DMN activity [26, 27] (whether it is related to the increased volume of pDMN in SZ seen in this study), and/or their linked networks in SZ. In addition to the possible effects of CPZ equivalents explored in current work, future studies should also investigate anti-cholinergic burden as another likely cause of medication-induced cognitive variance [28]. Visualizing these dynamic changes in spatial networks and their coupling could help identify the mechanisms by which these networks expand/shrink and how they are affected in SZ. In addition, one could also employ other metrics for dynamic spatial brain network coupling beyond simple volumetric measures, as well as adapt and combine with other techniques to identify spatiotemporal structures, such as dynamic mode decomposition where modes have intrinsic temporal behaviors [29]. Future research could investigate the impact of sliding window lengths on dynamic spatial brain networks, their coupling, and the observed significant effects of SZ. Our work highlights the importance of capturing spatiotemporal structure within and among networks and should be explored further to identify possible indicators of healthy and disordered brain function.

## COMPLIANCE WITH ETHICAL STANDARDS

This study analyzed anonymized data from existing repositories, all collected under the oversight of an IRB [4, 6, 7].

## ACKNOWLEDGMENTS

This work is funded in part by the NSF grant 2112455 and the NIH grant R01MH123610 to Dr. Vince D. Calhoun, and the NIH grant 5R01MH119251 to Dr. Armin Iraji. We would like to thank the members of TReNDS Center for valuable discussions.

